# Skeletal muscle lineage is dispensable for appendage regeneration in axolotl

**DOI:** 10.1101/2022.06.10.495631

**Authors:** Yan Hu, Xiangyu Pan, Yu Shi, Yuanhui Qiu, Liqun Wang, Prayag Murawala, Yanmei Liu, Wanjin Xing, Elly M Tanaka, Ji-Feng Fei

## Abstract

Regeneration of a complex appendage structure such as limb requires hierarchical coordination of multiple types of tissues. Muscles, as one of the major cell masses in limbs, have been reported recently to be critical to guide other tissue regeneration in planaria, but its function and relationship to other cells in vertebrate complex regeneration have been unclear. Here, we use *Pax7* mutant axolotls, in which the limb muscle is developmentally lost, to investigate limb regeneration in the absence of skeletal muscle. We find that the pattern of regenerated limbs is normal in *Pax7* mutants compared to the controls. Lack of muscles do not affect the proliferation of fibroblasts, another major population in limbs. Furthermore, using single cell RNA-sequencing, we demonstrate that the cell type composition in completely regenerated limbs in *Pax7* mutants is similar to that in the controls, except the lack of cell types in muscle lineage. Our study reveals skeletal muscle is not required for the guidance of complex tissue regeneration in axolotls, and provides new views of the tissue hierarchy in vertebrate appendage regeneration.

## Introduction

The limb is a complex organ contains multiple types of tissues and cells, such as skin, bone, cartilage, muscle, nerve, fibroblast and tendon. Reconstructing such a structure, that is generally incapable in mammals, is a technical challenging issue. One of the major obstacles is the lack of sufficient and systematic understanding of the interaction and the hierarchy relationship of the cell/tissue types involved in complex tissue regeneration. Axolotl is a tetrapod vertebrate that is able to regenerate diverse types of organs including the limb. Importantly, the cell/tissue composition and the structure of axolotl limb are similar to that in higher vertebrates, and the lost cell/tissue types and the structure could be precisely reproduced during regeneration, that very much resemble the original limb before amputation or injury ^1-5^.

Using various amphibian appendage regeneration models, previous studies showed that several types of cells and tissues, such as nerves, macrophages, apical epithelial cap and a recently identified regeneration-organizing cells (ROCs), play an essential role in mediating the injury responses or stimulating proliferation of many other cell types during complex tissue regeneration ^6-11^. Limbs fail to regenerate properly upon denervation. Without nerve tissue, the wound healing process seems normal, but blastema formation, in the other words, proliferation of the progenitors that participate in regeneration is impaired ^7, 10, 11^. Several studies identified that nerve stimulates progenitor activities through nerve-secreted factors such as BMP, NRG1 and MC4R ^12-14^. Furthermore, recent studies revealed that macrophages are also critical in successful complex tissue regeneration. Chemical depletion of macrophages leads to the failure of limb regeneration, but not the wound closure ^9, 11^. In addition, apical epithelial cap during salamander limb regeneration and ROCs in *Xenopus* tail regeneration, both derived from skin structure upon injury, have been reported to be one of the early-formed signaling centers to guide the underlying cells to form the blastema ^6, 8, 10, 11^. These works suggest that some tissue/cells, such as nerves, macrophages, apical epithelial cap and ROCs sit at higher hierarchy in regeneration events, either producing a regeneration-permissive environment or triggering directly progenitor proliferation, to guide the downstream regenerative responses of other cell types.

Muscle is one of the major cell types in appendages including limbs. In addition to its locomotor function, it has been reported that muscle secretes cytokines and peptides, and actively interact with other tissues ^15^. Furthermore, recent studies revealed that muscle plays critical roles in regulating regenerative responses in planaria, particularly providing the positional instructions to surrounding cell types and tissue, to maintain the proper regeneration ^16, 17^. In axolotl limb regeneration, connective tissues, but not the muscles follow the rule of distal transformation (giving rise to tissues only distal to the amputation plane), and differentially express the transcription factor MEIS (a positional identity regulator) along the proximo-distal axis ^18^. It suggests muscles do not harbor positional information to guide other tissue regeneration. However, direct evidences support this concept are still missing, and little is known about other potential roles of muscle, in the context of its relationship to other cell types, such as proliferation in complex tissue regeneration in vertebrate.

In this study, we take the advantage of a *Pax7* mutant axolotl line, in which the limb satellite cells and muscles are lost since early development ^19^, and test the role of muscles in limb regeneration. We find that in the absence of the limb muscle lineage, the patterning, early injury response and proliferation of major progenitors are proper in the process of limb regeneration. Single-cell RNA sequencing (scRNA-seq) reveals that the cell composition in regenerated limb of *Pax7* mutants is identical, except the loss of cells in muscle lineage, to the controls. Our work provides direct evidences that muscle lineage does not carry positional information and other instructions to regulate other tissues activity, but solely responsible to its own in complex tissue regeneration.

## Results

### The morphology and pattern of regenerated limb is proper in the absence of skeletal muscle

To test the role of muscle tissue in complex tissue regeneration, we chose previously reported *Pax7* mutant axolotls, in which the limb skeletal muscles and satellite cells are completely lost due to the early developmental defects (Supplementary Fig. S1) ^19^. When compared to controls, the overall size of limbs is slightly smaller in early juvenile *Pax7* mutants, due to the lack of limb muscle lineage. This discrepancy of the size of both limb and trunk increases with the proceeding of development, between *Pax7* mutants and the controls ^19^. It is likely a side effect of the massive loss of trunk muscles at later stages ^19^, which disable *Pax7* mutants to acquire sufficient food and nutrition to support their growth. However, except for the muscle lineage, all other major tissues, such as bones, cartilages, nerves and connective tissues are to some extent, normally developed in early juvenile *Pax7* mutants (Supplementary Fig. S2) ^19^. It allows us to investigate limb regeneration in the absence of muscle tissues. We first carried out limb amputation on both early juvenile healthy controls and muscle-free *Pax7* mutants, and followed the entire limb regeneration process. We found that the global morphology of blastema formation, growth and patterning of the limbs in *Pax7* mutants, based on the observation of over 30 individuals, are very similar to that in the controls (Fig. 1a).

**Figure 1.**
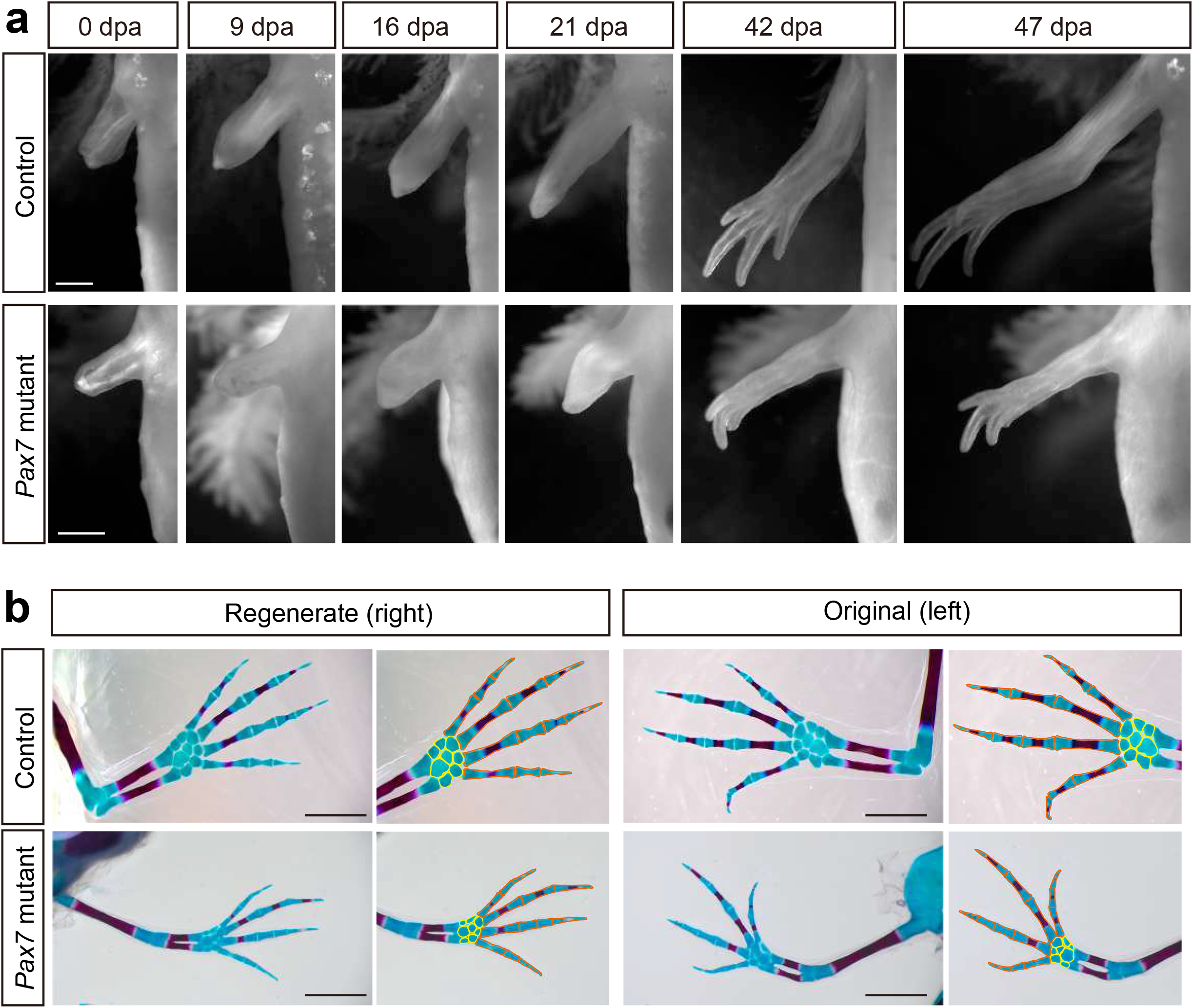
Morphology and patterning of limb regenerates in *Pax7* mutants. (a) Bright-field images of the limbs at indicated day post amputation (dpa), from 3-month-old controls and *Pax7* mutants. Scale bars, 1 mm. (b) Alcian blue and Alizarin red-stained fully regenerated (left panels) and untouched original (right panels) forelimbs from controls (upper panels) and *Pax7* mutants (lower panels). Amputations were carried out on 6-month-old animals, then allowed them to regenerate for 10-months for analysis. Hand part of each stained forelimb is presented at higher magnification, in which the red and yellow lines outline the phalange, metacarpal bones and carpal bones, respectively. Scale bars, 2 mm.

We further examined the patterning of regenerated limbs of the *Pax7* mutants and the controls in more detail. The bones and cartilages are generally used as an indication for the patterning of the limb regenerates ^20, 21^. Alcian blue/alizarin red staining revealed that only the bone and cartilage structures distal to the amputation plane are regenerated, and the newly formed phalange, metacarpal and carpal bones are properly patterned in *Pax7* mutants, which resembles the situation in the controls (Fig. 1b). Amputation on both early (only cartilages in forelimbs) and late stage (calcified bone in forelimbs) juvenile axolotls yield similar regeneration phenotypes (Supplementary Fig. S3 and Fig. 1b). We observed a slightly delayed regeneration (Fig. 1a) and bone calcification (Fig. 1b) in *Pax7* mutants compared to the controls. It is in line with the delayed limb developmental phenotype in late stage *Pax7* mutants ^19^. Presumably, the muscle defects result in the disability of mutants to eat enough Artemia food, therefore lack of enough nutrition to support regeneration. These results suggested that the skeletal muscles do not harbor the positional information and likely other crucial cues to guide the patterning in a complex tissue regeneration.

### Macrophage recruitments in early injury response do not require muscles

To figure out whether there are any changes on early injury responses in the absence of muscles, we chose the immune response, one of the early injury-induced events that has been reported to play essential role in limb regeneration ^9^, and examined the immune cell behavior in *Pax7* mutants. We carried out amputation on the upper limbs in both of the *Pax7* mutants and the controls, and collected samples at 0, 4, 9 and 16 dpa, which covers from early blastema to early differentiation stages. We used IBA1 as a marker to identify the macrophage cells on the limb longitudinal sections. The immunohistochemistry results showed that IBA1-positive macrophages are enriched to the amputation plane at early regeneration phase (4 dpa) in the controls, gradually declined at 9 dpa and finally returned back to about normal level at 16 dpa, as that observed at 0 dpa (Fig. 2a-c). Although the absolute number of IBA1-positive cells is reduced throughout the entire regeneration stages in *Pax7* mutants compared to the controls (Fig. 2b), when presenting as a percentage over all blastema cells, the proportion and dynamics of macrophages in limb regeneration of *Pax7* mutants resemble the situation in the controls (Fig 2c). This observation suggested that the muscles likely are not involved in the recruitment of macrophage at early regeneration stages.

**Figure 2.**
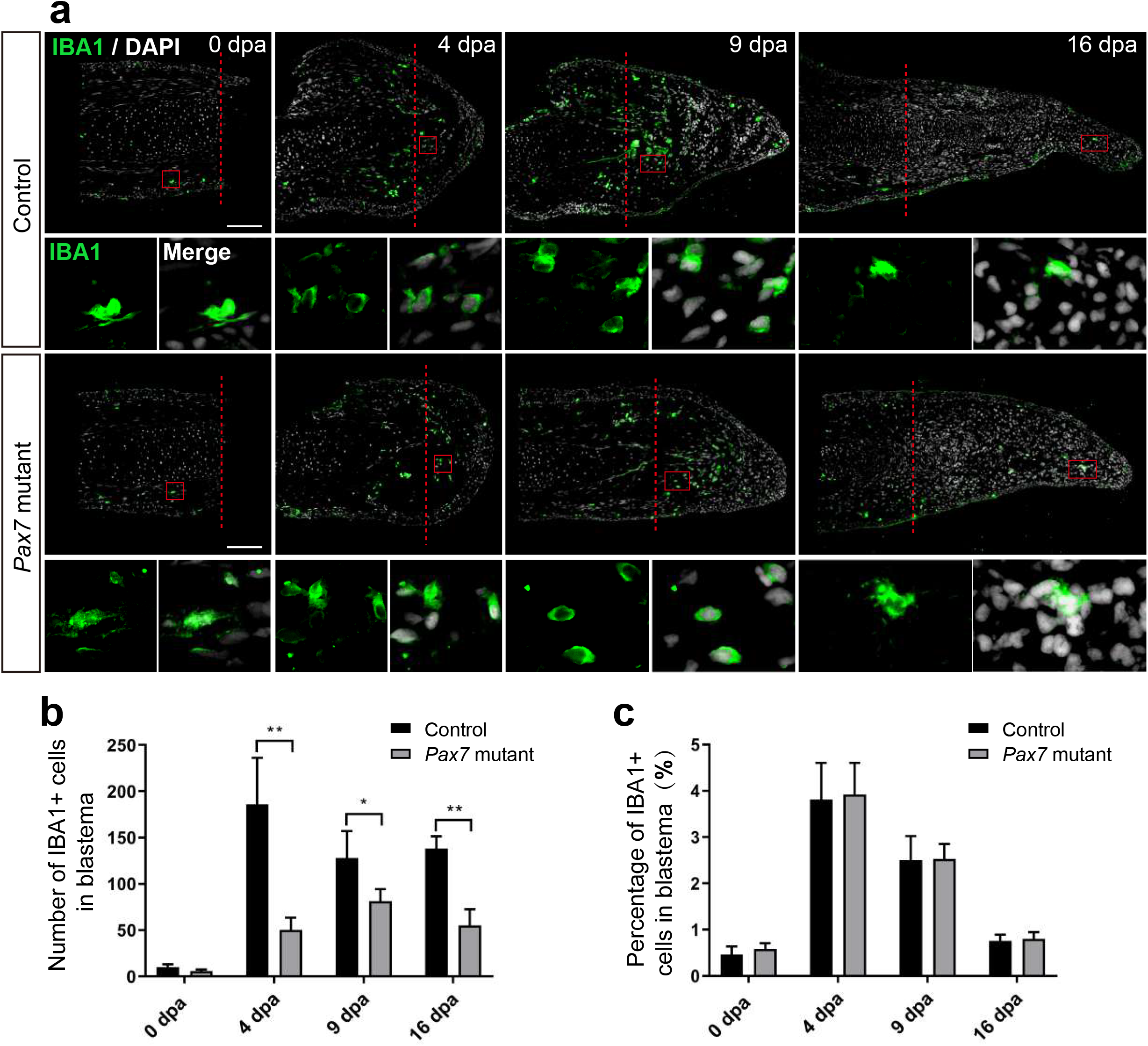
Macrophage dynamics in limb regeneration of *Pax7* mutants. (a) Immunofluorescence images of IBA1 (green) and DAPI (white) in limb longitudinal-sections, at 0-, 4-, 9- and 16-day post amputation (dpa), from control and *Pax7* mutant animals. Boxed regions are shown at higher magnification as separated channels. Read dashed lines, amputation planes. Scale bars, 200 μm. (b) and (c) Quantification of the absolute number (b) and the percentage (c) of IBA1 positive cells, within the areas of the newly formed blastema combined 500 μm zone proximal to the amputation plane, at 0-, 4-, 9- and 16-day post amputation (dpa), from controls (n=4) and *Pax7* mutants (n=4). Data are presented as mean ± sem, \**p* < 0.05, \*\**p* < 0.01.

### Fibroblast proliferation is not affected in the absence of skeletal muscle in limb regeneration

We next checked whether muscles play any role on the proliferation of blastema cells to guide the blastema formation. Previous studies have shown that limb blastema is the mixture of lineage-restricted progenitors ^22^. Connective tissue (fibroblast) is one of the major cell populations comprising the blastema, and has been reported to play a critical role in axolotl appendage regeneration ^23-27^. Therefore, we choose fibroblasts as a target for the examination of cell expansion and cell cycle dynamics in limb regeneration without muscles. We performed limb amputation and sample collection on the *Pax7* mutants and the controls as above described, additionally introducing a single pulse of EdU prior to sample collection. We then carried out immunohistochemistry using antibodies again PRRX1 (fibroblast) and Phospho-histone H3 (PH3, mitotic cells) on longitudinal sections, combined with EdU detection, to highlight fibroblast and reveal their dynamics (Fig. 3a). Immunohistochemistry and quantification of PRRX1-positive cells showed that the absolute number of PRRX1-positive fibroblasts is less in the *Pax7* mutants throughout the process of regeneration, when compared to the controls (Fig. 3a, 3b). It may represent the ground status of PRRX1-positive fibroblast reduction in *Pax7* mutants prior to limb amputation, since the size of the limb is smaller in *Pax7* mutants. However, with the proceeding of regeneration, the number of fibroblasts gradually increase in blastema in both *Pax7* mutants and controls. And the proportions of fibroblasts in blastema are comparable (slightly increased, but statistically not significant) at each regeneration time point in *Pax7* mutants, when compared to the controls (Fig. 3b, 3c), that is in line with the fact that myocytes and satellite cells comprise only a relatively small proportion of cells in early limb blastemas (Supplementary Fig. S4). Furthermore, EdU-positive or PH3-positive cells, which label fibroblasts at S- or M-phases of the cell cycle respectively, also occupied similar ratio in the *Pax7* mutants and controls at all regeneration stages analyzed (Fig. 3c, 3d). These results indicated cell cycle dynamics of PRRX1-positive fibroblasts are not dramatically affected in the absence of muscles during limb regeneration.

**Figure 3.**
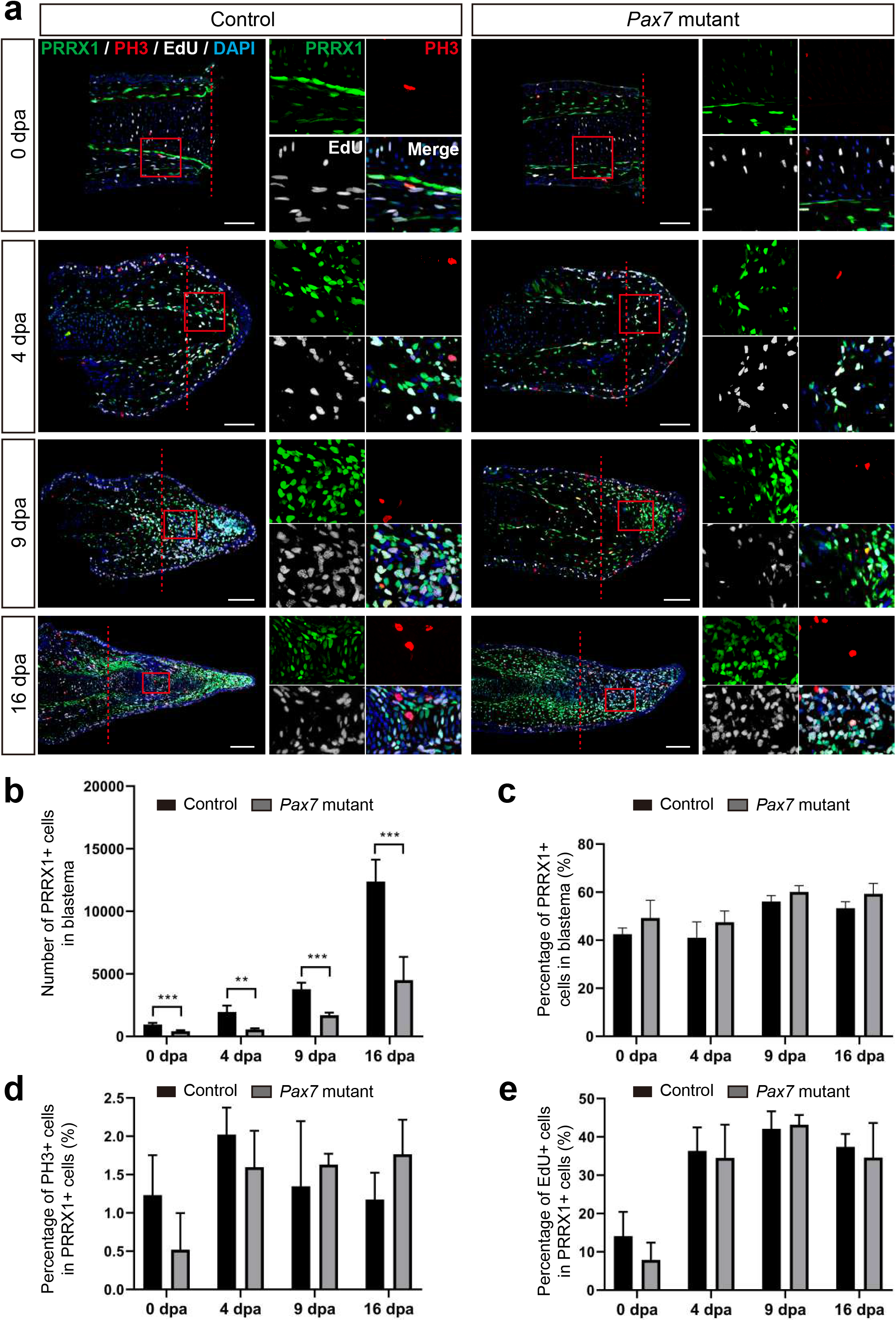
Fibroblast dynamics during limb regeneration in the absence of muscles. (a) Immunofluorescence images of PRRX1 (green), PH3 (red), combined with EdU (white) and DAPI (blue) staining in limb longitudinal-sections, at 0-, 4-, 9- and 16-day post amputation (dpa), from control and *Pax7* mutant animals. Boxed regions are shown at higher magnification as separated channels. Read dashed lines, amputation planes. Scale bars, 200 μm. (b) and (c) Quantification of the absolute number (b) and the percentage (c) of PRRX1 positive cells, in the areas of the newly formed blastema combined 500 μm zone proximal to the amputation plane, at 0-, 4-, 9- and 16-day post amputation (dpa), from controls (n=4) and *Pax7* mutants (n=4). Data are presented as mean ± sem, \*\**p* < 0.01, \*\*\**p* < 0.001. (d) and (e) Quantification of the percentage of PH3- (d) and EdU- (e) positive cells, in PRRX1-positive cells quantified in (b).

### Full spectrum of cell types, except muscle lineage are regenerated in Pax7 mutant limbs

To determine whether other relevant cell types, in addition to fibroblasts, respond properly during regeneration in the absence of muscles, we systematically examined cell type composition in fully regenerated forelimbs from both *Pax7* mutants and controls. To this end, we collected uninjured and 47-days regenerated forelimbs from *Pax7* mutants and controls, isolated single cells and carried out scRNA-seq, which allowing unbiased identification of cell types (Fig. 4a). In total, we obtained 87,177 high-quality cells for single cell transcriptome and cell type identity analysis. After clustering all cells and reflecting them in uniform manifold approximation and projection (UMAP), we identified 41 putative cell populations. We then used typical cell type markers identified previously to assign each cluster ^23-25, 28, 29^. The finally defined clusters belong to 13 cell types which cover all major limb cell types (Fig. 4b), such as fibroblastic connective tissue (CT) cells, macrophages, Schwann cells, and chondrocytes. We found that in the uninjured limbs, except satellite and myocytes, all other cell types are identified in both *Pax7* mutants and controls (Fig. 4b, 4c). Moreover, cell type composition in the 47-days regenerated limbs recapitulates the situation in uninjured limbs, from which only the cell types in muscle lineage abruptly decline in the *Pax7* mutant limb regenerates (Fig. 4b, 4c). Meanwhile, we also observed that the proportions of the majority of cell types are nearly unchanged in the *Pax7* mutants compared to the controls, except for the decreased epidermal cell and increased skeletal myocyte levels in the control regenerates (Fig. 4b, 4c). We chose several major marker genes of all the 13 cell types, and analyzed the expression of these genes in all major identified cell types in both *Pax7* mutants and controls. We found that cells in *Pax7* mutants largely share common gene expression patterns with the cells in controls (Fig. 4d). Our data suggest that the deficiency of limb muscles do not change the cell identities and states at a systematic molecular level. Moreover, we performed immunohistochemistry on cross sections of 47-days limb regenerates from both *Pax7* mutants and controls, using antibodies against MHC, PRRX1, MBP, βIII-tublin and SOX9, which labels muscle cells, fibroblastic CT cells, Schwann cells, neuronal cells and chondrocytes, respectively. We found that, in consistent with the scRNA-seq data, all these identified cell types are all present in the regenerated limb in *Pax7* mutants. And the distribution of each cell type in limb regenerates resembles its location in original uninjured limb (Fig. 5a-c). These results reveal that all other cell types in limb regenerate properly without the support of muscle tissues. Therefore, we conclude that muscles do not play essential roles on the regeneration of other cells types in complex tissue regeneration in axolotls.

**Figure 4.**
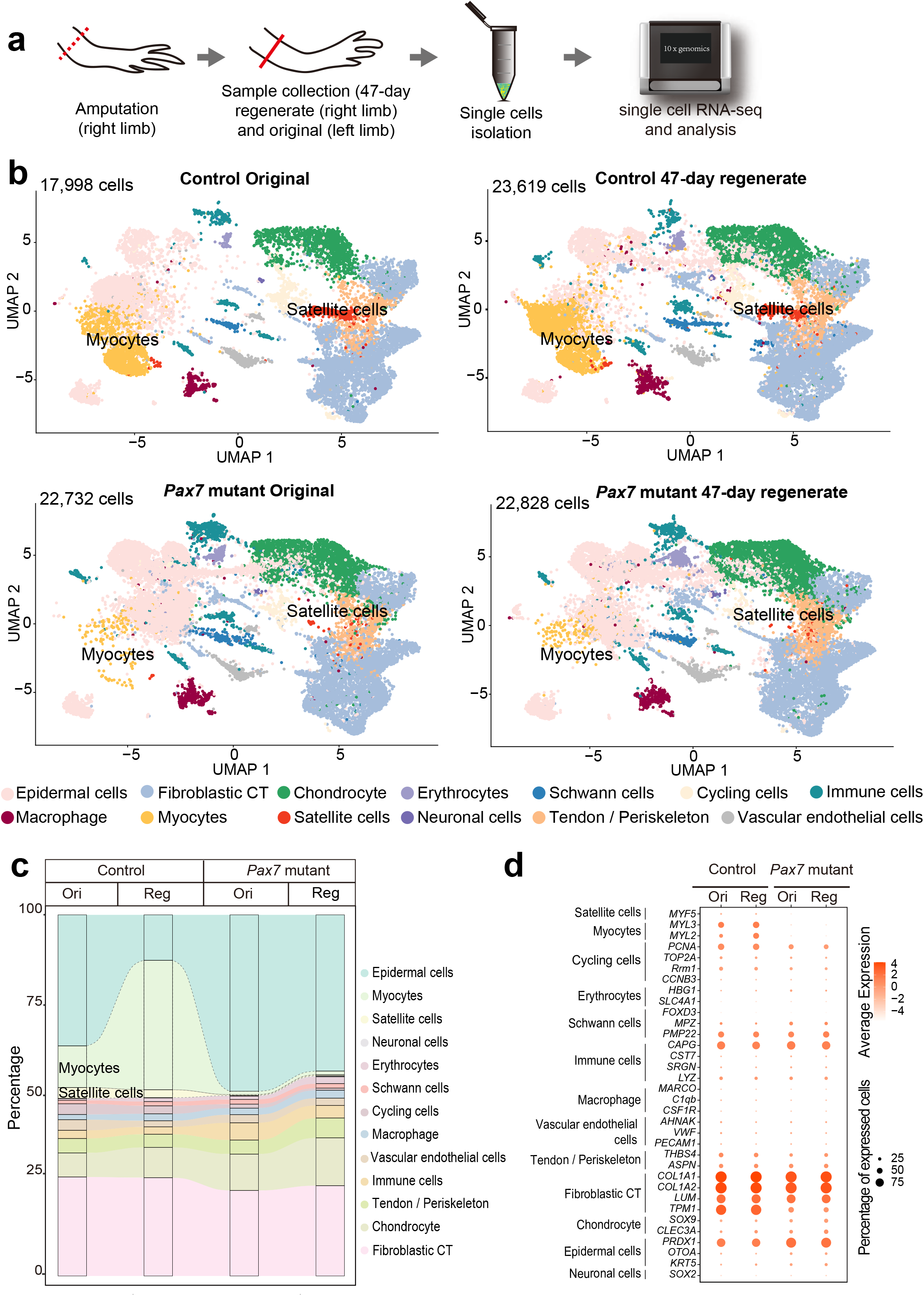
Integrated transcriptional cell state atlas of the axolotl limb from the 47-day regenerate and original in both controls and *Pax7* mutants. (a) Scheme of the scRNA-seq workflow used for the limb tissues collected from the 47-day regenerate (right limb) and original (left limb) in both controls and *Pax7* mutants. Dotted line, amputation plane; line, the position of sample collection. (b) UMAP visualization of the scRNA-seq data from four sampling stages. The cell-type annotation is determined by published cell-lineage specific markers. Two muscle-related cell types are highlighted. (c) Sankey plot showing the percentage of all 13 cell types from the 47-day regenerate and original limbs in both controls and *Pax7* mutants. (d) Dot plot visualizing the expression of marker genes of all 13 cell types shared between the controls and *Pax7* mutants. The circle size represents the percentage of cells at each sample expressing the gene, and color represents the average expression level.

**Figure 5.**
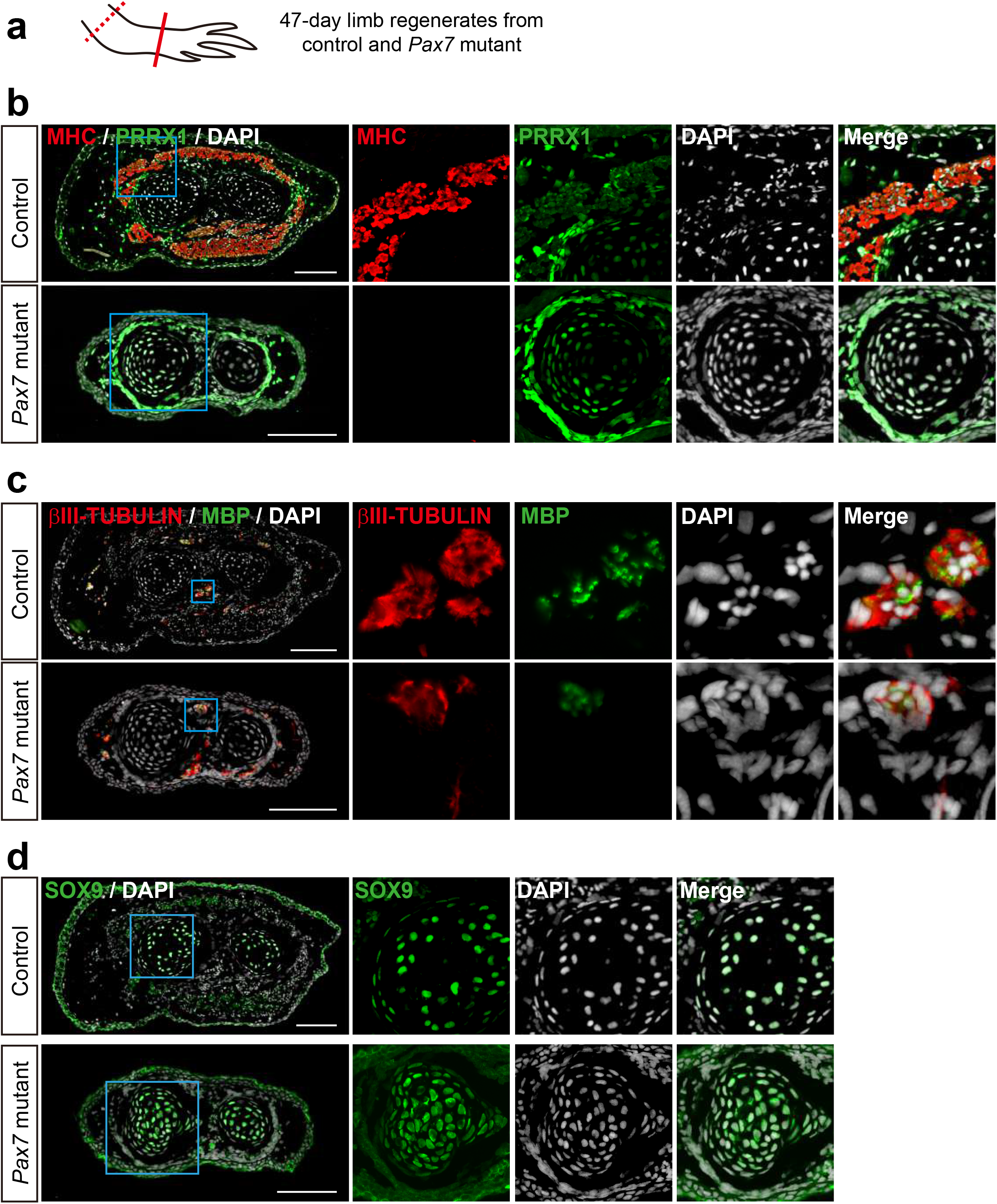
Examination of cell types in limb regenerates in *Pax7* mutants. (a) Scheme of the sample collection. Dotted line, amputation plane; line, the position of analyzed section. (b-d) Immunofluorescence images of MHC (red), PRRX1 (green) and DAPI (white) (b); βIII-TUBULIN (red), MBP (green) and DAPI (white) (c); and SOX9 (green) and DAPI (d), in cross-sections, from 47-days fully regenerated control and *Pax7* mutant limbs. Boxed regions are shown at higher magnification as separated channels. Scale bars, 200 μm.

## Discussion

Although the function of muscles has been studied during the regeneration process of planarians ^16, 17^, the potential roles of muscles involved in the hierarchical coordination of limb regeneration of vertebrates have not been comprehensively explored. Here, we used a stable germline transmitted *Pax7* mutant axolotl line, in which the loss of limb muscle phenotype resembles the situation in *Pax3* mutants in other species (loss of limb muscle) ^19^, to test the role of muscle tissue in complex tissue regeneration. By combining immunostaining and scRNA-seq approaches of control and *Pax7* mutant axolotl lines to survey molecular activities of the role of muscles during limb regeneration, we found that muscle lineage is dispensable for the patterning and dynamics of other cell types in tissue regeneration.

### The role of muscle in hierarchical pattern formation during axolotl limb regeneration

The mature limb consists of multiple tissues, including the epidermis, dermis, muscle, nerve, blood vessels and skeletal elements. These tissues must regenerate coordinately according to the patterning hierarchy to restore functionality ^22, 30^. Previous studies have found that some cell types, such as nerve and fibroblast CT cells are key for axolotl limb regeneration ^24-26, 31^. As one of the major cell masses in limbs, in addition to supporting the skeleton and controlling movement, other roles during homeostasis and regeneration are not well established. In newts, another salamander species, *Pax3* mutation causes the loss of limb muscle during development. Investigation on the limb regeneration using F0 *Pax3* mutant newts revealed that those animals can regenerate the limb after amputation ^32^. However, a detailed characterization of the patterning and the dynamics of other cell types in muscle-free limb regeneration was still missing. Here, our multi-dimension studies reveal that lack of muscles do not affect the limb regeneration in axolotls. Firstly, the experimental data showed that muscles do not retain positional memory and can be viewed as being positionally naive. According to the defined cell types within the patterning hierarchy model (pattern forming cells and pattern following cells) ^33^, muscle should be the pattern following cells in axolotl. Secondly, muscle tissues do not play a dominant role on regulating the early injury responses or proliferation of other cell types during regeneration. Overall, our data indicate that muscles locate at the bottom of the hierarchical pyramid in complex tissue regeneration.

### The role of muscle during evolution

In planaria, muscles provide cues of positional information to guarantee the precision of the regeneration ^16, 17^. Nacu and colleagues’ work suggests that muscles may function differently in axolotls. It is very likely muscles do not harbor positional information, instead connective tissues may be the source of the positional instructions ^18^. Here, we demonstrate that the patterning of limb regenerates is not affected in the absence of limb skeletal muscles, and provide direct evidence that the muscle lineage does not give positional instructions, as well as other major instructions to guide surrounding cell activities in a complex tissue regeneration. Interestingly, a recent study showed that muscles in planaria function to some extent as connective tissues ^34^. It is reasonable to assume that during evolution, the divergence of muscles gives rise to the connective tissues, and part of the functions originally undertaken by muscles are segregated and re-assigned to newly derived connective tissues (fibroblasts). It will be necessary to directly test the role of fibroblasts in vertebrates on positional determination in future, such as using genetic approach to ablate the population of fibroblasts, then follow the regeneration of complex tissue in the absence of connective tissues.

### The lineage switch of muscle

The regenerative environment has the potential to induce differentiation, de-differentiation or trans-differentiation of particular cell types. A classical trans-differentiation observation in regeneration is reported in the lens regeneration in newts ^35^. When the lens is removed, dorsal iris pigmented epithelial cells could trans-differentiate into lens tissue ^35^. In terms of muscle tissue, several recent studies have revealed that fibroblasts are capable of trans-differentiating into skeletal muscles *in vitro* under given conditions ^36 37^. In our work, we found that, although all other cell types, except the muscle lineage, are present in the limb, upon amputation, the regeneration environment itself is not sufficient to trigger the trans-differentiation of other lineage into skeletal muscles.

Overall, our study uncovered the dispensable role of muscle during axolotl limb regeneration, covering the activities of macrophages immune response, fibroblast proliferation, and other cell types hierarchical coordination to reconstruct the limb. It gives new insights into exploring essential components of tissue regenerating scaffolds, ranging from the macroscopic to the molecular scale in a hierarchical manner. Moreover, our finding may contribute to the realization of regenerative therapies by building biomaterial constructs *in vitro*.

## Materials and methods

### Axolotl care and operation

Animal experiments were carried out according to the guidelines of the ethics committee of Guangdong Provincial People’s Hospital. In this study, we used d/d and *Pax7*-CRIPSR mutant, strain tm(*Pax7*^*4751V6D20/4754V6D20*^)^LABF^ axolotls (*Ambystoma Mexicanum*) ^38^, aging between 2.5 and 16 months for analysis. Axolotl larvae were kept individually in plastic containers with a change of fresh tap water once a day and fed artemia daily. Axolotls were anaesthetized with 0.01% ethyl-p-aminobenzoate (Benzocaine; Sigma) solution prior to amputation, imaging, EdU injection and sample collection. Amputation was carried out on right forelimb of axolotl larvae (about 3-month-old, except for one bone regeneration experiment, 6-month-old animals were used) by cutting at the mid-upper arm followed by trimming of the bone to allow proper blastema formation. Bright-field axolotl limb images were obtained using an Olympus stereomicroscope. Samples of uninjured limbs or limb regenerates were collected at the stages indicated for analysis. If applicable, a single pulse of EdU (at a dose of 10 mg/kg body weight) was introduced into axolotls by intraperitoneal injection, and kept for 3 hours prior to sample collection. All samples collected were fixed in 1 × MEMFA (0.1M MOPS, pH 7.4, 2mM EGTA, 1mM MgSO_4_ and 3.7% formaldehyde) for further analysis, unless specified.

### Alcian blue and Alizarin red staining

Samples were fixed in 4% PFA (4% paraformaldehyde prepared in phosphate buffer, pH 7.2) for at least 24 hours for Alcian blue and Alizarin red staining. Briefly, after several washes in PBS, the internal organs of the fixed axolotls were carefully removed, then sequentially dehydrated in 25%, 50% and 70% ethanol solutions. Firstly, the samples were stained for cartilage tissue with 0.01% Alcian blue 8GX solution (prepared in ethanol and glacial acetic acid, 3:2) for approximately 48 hours, then gradually rehydrated with 70%, 50% and 25% ethanol series, washed with distilled water for 24-48 hours, followed by an enzymatically digested in 1% trypsin (MP Biomedicals, 153571) for 1-2 hours at 37 °C. After digestion, samples were rinsed in 1% KOH, then stained for bone tissue with 0.01% Alizarin Red solution (prepared in 1% KOH) overnight, followed by a series of 1% KOH washes until the samples look transparent. lastly, the samples were dehydrated in 25%, 50%, 70%, 90% and pure ethanol series, then incubated sequentially in glycerol/ethanol series (1:3, 1:1, 3:1), and finally stored in glycerol at room temperature. Images were acquired with an Olympus SZX10 microscope (Olympus, Tokyo, Japan).

### Immunohistochemistry and EdU staining

We collected 10 μm-thick cryosections in accordance with standard procedure for Immunohistochemistry and EdU staining. When combined with EdU detection, we carried out firstly EdU staining using Click-iT EdU kit (Invitrogen, C10340) in accordance with the manufacturer’s instructions, then followed by a standard immunohistochemistry protocol as previously described ^39^. In brief, after PBS wash and permeabilization with PBST, slides were blocked in 5% serum prepared in PBST, then incubated with primary antibodies overnight at 4 °C, followed by sequential PBST washes and secondary antibody (with DAPI, Sigma-Aldrich, D9542) incubation, and finally mounted using Mowiol 4-88 (Sigma-Aldrich, 9002-89-5) mounting medium after several PBST washes. Fluorescence images were acquired with an Olympus IX83 microscope (Olympus, Tokyo, Japan), using 20× objectives and analyzed using FIJI. The following antibodies were used in this study, MBP (Genetex, GTX761141), PRRX1 ^25^, βIII-TUBULIN (R&D, MAB1195), SOX9 (Chemicon, Ab5535), PAX7 (DSHB, AB528428), MEF2C (Santa Cruz, sc-365862), phospho-Histone H3 (Abcam, 10543), MHC (DSHB, A4.1025). Alexa 488-donkey-anti-mouse IgG (Jackson, 711-547-003), Alexa 555-donkey-anti-rat IgG (Invitrogen, SA5-10027), Alexa 647-donkey-anti-mouse IgG (Jackson, 715-607-003), CY3-donkey-anti-mouse IgG (Jackson, 715-165-151).

### Single cell dissociation of scRNA-seq

For each scRNA-seq experiment, 30-40 original or regenerating forelimbs were collected as a pool for single cell dissociation. Each sample was minced and digested in 1000 μL 1 × Liberase (Roche) diluted in 0.8 × PBS- (without Mg^2+^/Ca^2+^) supplemented with 0.5 U/μL Dnase I (Thermo Scientific) at room temperature for approximately 30 minutes. The reaction was stopped by adding 10% FBS in PBS. Then, the cell suspension was filtered through a 70 μm cell strainer, followed by a centrifugation at 500 g for 5 minutes. Cells were resuspended in 2 mL 0.8 × PBS and filtered through a 30 μm cell strainer, followed by a centrifugation at 500 g for 5 minutes. Finally, the cells collected were resuspended in 0.5 mL 0.8 × PBS to generate a single-cell suspension used for the library construction.

### Library preparation and sequencing

The scRNA-seq library was constructed using the Chromium single-cell 3 prime v2 reagent kit (10× Genomics) in accordance with the manufacturer’s instructions (https://support.10xgenomics.com/single-cell-gene-expression/index/doc/user-guide-chromium-single-cell-3-reagent-kits-user-guide-v2-chemistry). All libraries were further prepared based on the requirements of the Nova-seq6000 sequencing platform manufactured by MGI®. The DNA concentration was determined by a Qubit (Invitrogen®). Then, samples with 2 pmol of nucleotides were pooled to generate single-strand DNA circles (ssDNA circles). DNA nanoballs (DNBs) were generated with the ssDNA circles by rolling the circles during replication to significantly increase the fluorescent signals during the sequencing process. The DNBs were loaded into the patterned nanoarrays and sequenced on the Nova-seq6000 sequencing platform with a paired end read length of 28-150 bp.

### Single-cell RNA-seq data processing

We firstly processed 10× Genomics raw data using the Cell Ranger Single-Cell Software Suite (version 4.0.0), including using ‘cellranger mkfastq’ to demultiplex raw base call files into FASTQ files, then using ‘cellranger count’ to perform alignment, filtering, barcode counting, and UMI counting. The reads were aligned to the axolotl reference genome AmbMex60DD ^40^ (https://www.axolotl-omics.org/assemblies) with STAR ^41^ which was imbedded in the 10× Genomics Cell Ranger-4.0.0 software with default parameters to generate the absolute Unique Molecular Identifier (UMI) counts. The output from different lanes was finally aggregated using ‘cellranger aggr’ with the default parameter setting. Then we mapped UMIs to genes, followed by removing low-quality cells. Cells with fewer 200 unique genes were excluded for further analysis. Then we utilized functions in the Seurat package to normalize and scale the single-cell gene expression data. The data were first normalized using the ‘NormalizeData’ function with the normalization method set to ‘LogNormalize’. Specifically, the expression of each gene *i* in cell *j* was determined by the UMI count of gene *i* divided by the total number of UMI of cell *j*, followed by multiplying by 10000; for the normalization the log-transformed counts were then computed with a base of 2. We then removed the irrelevant sources of variation by regressing out cell-cell variation within gene expression driven by batch and the number of detected UMI; this was implemented using the ‘ScaleData’ function. Finally, the corrected expression matrix was used as an input for further analysis.

### Dimension reduction, cell clustering

To enhance the identification of common cell types and enable comparative analyses of different experiments, we integrated datasets from all stages by using the ‘FindIntegrationAnchors’ function in Seurat, which was implemented in the Seurat workflow (https://satijalab.org/seurat/v3.0/immune_alignment.html).

We next restricted the corrected expression matrix to the subsets of highly variable genes (HVGs), and then centered and scaled values before performing dimension reduction and clustering on them. Methodologically, the top 2000 HVGs in single-cell data were selected by first fitting a generalized linear model to the mean-dependent trend for the gene-specific variance of all genes, and selecting genes that deviated significantly from the fitted curve. This was implemented using the ‘FindVariableGenes’ function in the Seurat package by setting the valid value of average expression as a range from 0.05 to 5 and that of dispersion as no less than 0.5.

We then used the ‘RunPCA’ function in the Seurat package to perform principal component analysis (PCA) on the single-cell expression matrix with genes restricted to HVGs. Given that many principal components explain very low proportion of the variance, the signal-to-noise ratio can be improved substantially by selecting a subset of significant principal components. The number of significant principal components was determined by the permutation test, implemented using the permutationPA function from the jackstraw R package (https://cran.r-project.org/web/packages/jackstraw). We then utilized the ‘FindClusters’ function in the Seurat package to conduct the cell clustering analysis through embedding cells into a graph structure in PCA space. Due to the large number of cells in our study, we set the parameter resolution to 0.8. This identified a total of 41 clusters. The clustering results were presented using a uniform manifold approximation and projection (UMAP) based on plots generated by the ‘RunUMAP’ function. The ‘FindAllMarkers’ function was used to identify differentially expressed genes with default parameters. Finally, top 100 differentially expressed genes in each cluster were kept as marker genes of each cluster.

### Identification of cell types

To obtain the comprehensive genetic annotation information, we reannotated the gene annotation of axolotl using the protein sequence. The human (GRCh38) and mouse (GRCm38) protein sequences were downloaded from the NCBI. We then aligned all the public data against the axolotl protein sequence by using all *vs*. all Blast ^42^ (version 2.2.25) with the main parameter of “–e 1e-5”. The best hit result from alignments with a Z-score ≥ 200 was used to perform the homologous annotation. Then, top 100 marker genes in each cluster were annotated manually by querying literatures one by one to define the cell types.

### Quantifications, and Statistical analyses

All quantifications were carried out manually. For each sample, we collected a series of 10 μm longitudinal limb sections for immunohistochemistry and quantification. The adjacent sections have an 80 μm interval. Generally, each control and *Pax7* mutant sample contains 11-14 and 7-9 sections, respectively. All relevant cells from this series of sections in the region of interest were summed for statistics. All immune- or DAPI-positive cells in newly regenerated blastema and within 500 μm zone proximal to the amputation plane were used for quantification. All data are presented as mean ± sem, an unpaired t-test was used to determine whether differences in the experimental group were statistically significant. Student’s t-tests were performed using Microsoft Excel and Graphpad Prism 9.0.0. A p-value less than 0.05 was considered to be statistically significant.

## Acknowledgments

This study was supported by the National Key R&D Program of China (2019YFE0106700, 2021YFA0805000), the Natural Science Foundation of China (31970782, 32070819), the High-level Hospital Construction Project of GDPH (DFJHBF202103, KJ012021012), Project of Department of Education of Guangdong Province (2018KZDXM027), Key-Area Research and Development Program of Guangdong Province (2018B030332001) and Guangdong-Hong Kong-Macao-Joint Laboratory Program (2019B121205005).

## References

1. Brockes, J. P. & Kumar, A. Appendage regeneration in adult vertebrates and implications for regenerative medicine. Science 310, 1919–1923 (2005).

2. Haas, B. J. & Whited, J. L. Advances in Decoding Axolotl Limb Regeneration. Trends Genet. 33, 553–565 (2017).

3. Sessions, S. K., Gardiner, D. M. & Bryant, S. V. Compatible limb patterning mechanisms in urodeles and anurans. Dev. Biol. 131, 294–301 (1989).

4. Nacu, E. & Tanaka, E. M. Limb regeneration: a new development? Annu. Rev. Cell Dev. Biol. 27, 409–440 (2011).

5. Flowers, G. P., Sanor, L. D. & Crews, C. M. Lineage tracing of genome-edited alleles reveals high fidelity axolotl limb regeneration. Elife 6, (2017).

6. Aztekin, C., Hiscock, T. W., Marioni, J. C., Gurdon, J. B., Simons, B. D. & Jullien, J. Identification of a regeneration-organizing cell in the Xenopus tail. Science 364, 653–658 (2019).

7. Todd, T. J. On the process of reproduction of the members of the aquatic salamander. Quart. J. Scl. Lit. Arts 16, 84–96 (1823).

8. Thornton, C. S. Influence of an eccentric epidermal cap on limb regeneration in Amblystoma larvae. Dev. Biol. 2, 551–569 (1960).

9. Godwin, J. W., Pinto, A. R. & Rosenthal, N. A. Macrophages are required for adult salamander limb regeneration. Proc. Natl. Acad. Sci. U S A. 110, 9415–9420 (2013).

10. Makanae, A. & Satoh, A. Early regulation of axolotl limb regeneration. Anat. Rec. (Hoboken) 295, 1566–1574 (2012).

11. Stocum, D. L. Mechanisms of urodele limb regeneration. Regeneration 4, 159–200 (2017).

12. Satoh, A., Cummings, G. M., Bryant, S. V. & Gardiner, D. M. Neurotrophic regulation of fibroblast dedifferentiation during limb skeletal regeneration in the axolotl (Ambystoma mexicanum). Dev. Biol. 337, 444–457 (2010).

13. Farkas, J. E., Freitas, P. D., Bryant, D. M., Whited, J. L. & Monaghan, J. R. Neuregulin-1 signaling is essential for nerve-dependent axolotl limb regeneration. Development 143, 2724–2731 (2016).

14. Zhang, M. et al. Melanocortin Receptor 4 Signaling Regulates Vertebrate Limb Regeneration. Dev. Cell 46, 397–409 e395 (2018).

15. Pedersen, B. K. Muscle as a secretory organ. Compr. Physiol. 3, 1337–1362 (2013).

16. Witchley, J. N., Mayer, M., Wagner, D. E., Owen, J. H. & Reddien, P. W. Muscle cells provide instructions for planarian regeneration. Cell Rep. 4, 633–641 (2013).

17. Scimone, M. L., Cote, L. E. & Reddien, P. W. Orthogonal muscle fibres have different instructive roles in planarian regeneration. Nature 551, 623–628 (2017).

18. Nacu, E. et al. Connective tissue cells, but not muscle cells, are involved in establishing the proximo-distal outcome of limb regeneration in the axolotl. Development 140, 513–518 (2013).

19. Nowoshilow, S. et al. The axolotl genome and the evolution of key tissue formation regulators. Nature 554, 50–55 (2018).

20. Maden, M. Vitamin A and pattern formation in the regenerating limb. Nature 295, 672–675 (1982).

21. Thoms, S. D. & Stocum, D. L. Retinoic acid-induced pattern duplication in regenerating urodele limbs. Dev. Biol. 103, 319–328 (1984).

22. Kragl, M. et al. Cells keep a memory of their tissue origin during axolotl limb regeneration. Nature 460, 60–65 (2009).

23. Leigh, N. D. et al. Transcriptomic landscape of the blastema niche in regenerating adult axolotl limbs at single-cell resolution. Nat. Commun. 9, 5153 (2018).

24. Lin, T. Y. et al. Fibroblast dedifferentiation as a determinant of successful regeneration. Dev. Cell 56, 1541–1551 e1546 (2021).

25. Gerber, T. et al. Single-cell analysis uncovers convergence of cell identities during axolotl limb regeneration. Science 362, (2018).

26. McCusker, C. D. & Gardiner, D. M. Positional information is reprogrammed in blastema cells of the regenerating limb of the axolotl (Ambystoma mexicanum). PloS one 8, e77064 (2013).

27. Currie, J. D., Kawaguchi, A., Traspas, R. M., Schuez, M., Chara, O. & Tanaka, E. M. Live Imaging of Axolotl Digit Regeneration Reveals Spatiotemporal Choreography of Diverse Connective Tissue Progenitor Pools. Dev. Cell 39, 411–423 (2016).

28. Qin, T. et al. Single-cell RNA-seq reveals novel mitochondria-related musculoskeletal cell populations during adult axolotl limb regeneration process. Cell Death Differ. 28, 1110–1125 (2021).

29. Li, H. et al. Dynamic cell transition and immune response landscapes of axolotl limb regeneration revealed by single-cell analysis. Protein Cell 12, 57–66 (2021).

30. Aztekin, C. Appendage regeneration is context dependent at the cellular level. Open Biol. 11, 210126 (2021).

31. Wells, K. M., Kelley, K., Baumel, M., Vieira, W. A. & McCusker, C. D. Neural control of growth and size in the axolotl limb regenerate. Elife 10, (2021).

32. Elewa, A. et al. Reading and editing the Pleurodeles waltl genome reveals novel features of tetrapod regeneration. Nat. Commun. 8, 2286 (2017).

33. Vieira, W. A. & McCusker, C. D. Hierarchical pattern formation during amphibian limb regeneration. Biosystems 183, 103989 (2019).

34. Cote, L. E., Simental, E. & Reddien, P. W. Muscle functions as a connective tissue and source of extracellular matrix in planarians. Nat. Commun. 10, 1592 (2019).

35. Vergara, M. N., Tsissios, G. & Del Rio-Tsonis, K. Lens regeneration: a historical perspective. Int. J. Dev. Biol. 62, 351–361 (2018).

36. Boularaoui, S. M. et al. Efficient transdifferentiation of human dermal fibroblasts into skeletal muscle. J. Tissue Eng. Regen. Med. 12, e918–e936 (2018).

37. Xu, B., Siehr, A. & Shen, W. Functional skeletal muscle constructs from transdifferentiated human fibroblasts. Sci. Rep. 10, 22047 (2020).

38. Nowoshilow, S., Fei, J. F., Voss, R. S., Tanaka, E. M. & Murawala, P. Gene and transgenics nomenclature for the laboratory axolotl - Ambystoma mexicanum. Dev. Dyn., doi: 10.1002/dvdy.1351. Online ahead of print (2021).

39. Fei, J. F., Schuez, M., Knapp, D., Taniguchi, Y., Drechsel, D. N. & Tanaka, E. M. Efficient gene knockin in axolotl and its use to test the role of satellite cells in limb regeneration. Proc. Natl. Acad. Sci. USA 114, 12501–12506 (2017).

40. Schloissnig, S. et al. The giant axolotl genome uncovers the evolution, scaling, and transcriptional control of complex gene loci. Proc. Natl. Acad. Sci. USA 118, (2021).

41. Dobin, A. et al. STAR: ultrafast universal RNA-seq aligner. Bioinformatics (Oxford, England) 29, 15–21 (2013).

42. Altschul, S. F., Gish, W., Miller, W., Myers, E. W. & Lipman, D. J. Basic local alignment search tool. J. Mol. Biol. 215, 403–410 (1990).

