## Supplementary figure and legends for "Skeletal muscle lineage is dispensable for appendage regeneration in axolotl"

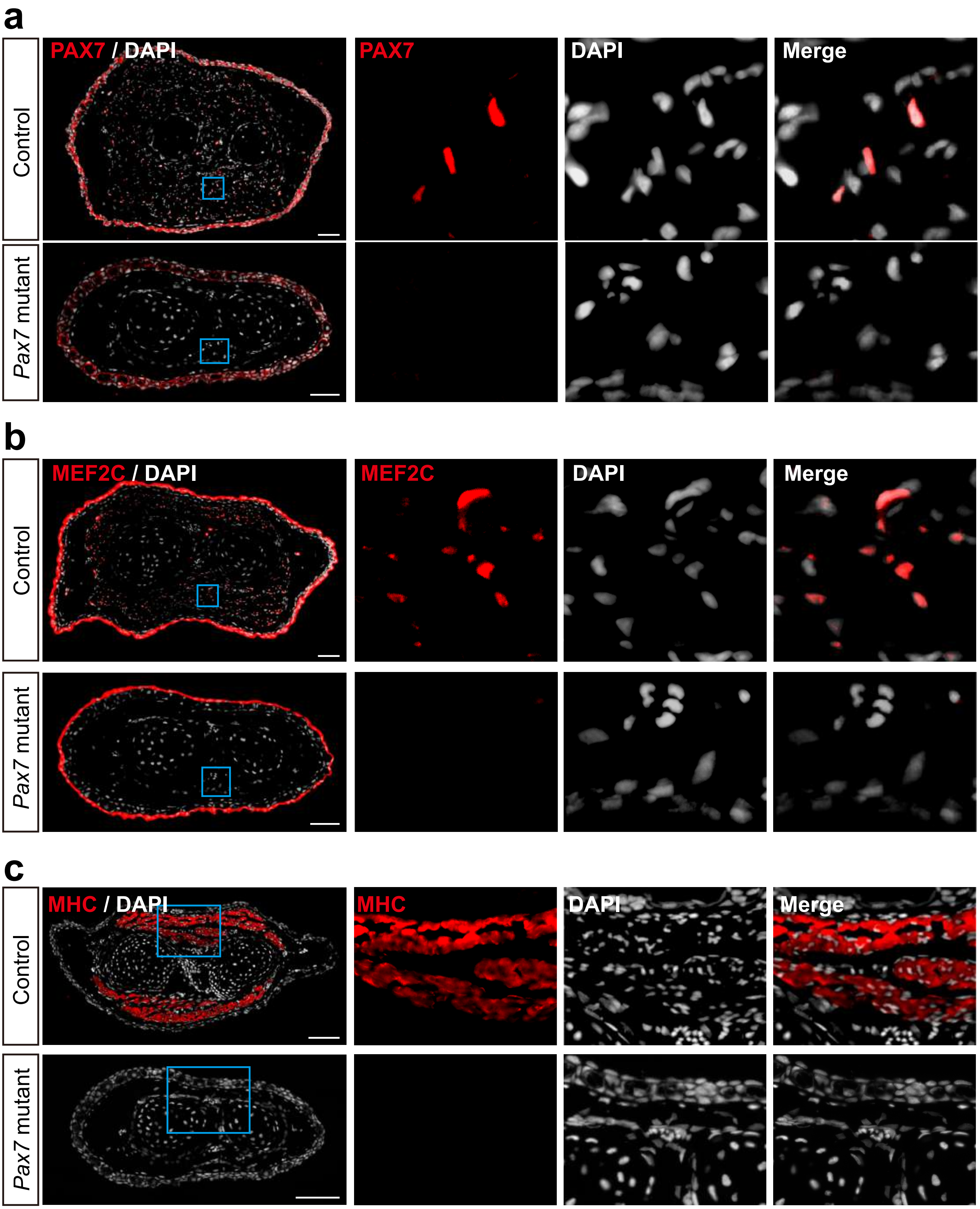

a
Control
Pax7 mutant
PAX7 / DAPI
PAX7
DAPI
Merge
b
Control
Pax7 mutant
MEF2C / DAPI
MEF2C
DAPI
Merge
c
Control
Pax7 mutant
MHC / DAPI
MHC
DAPI
Merge


Fig. S1

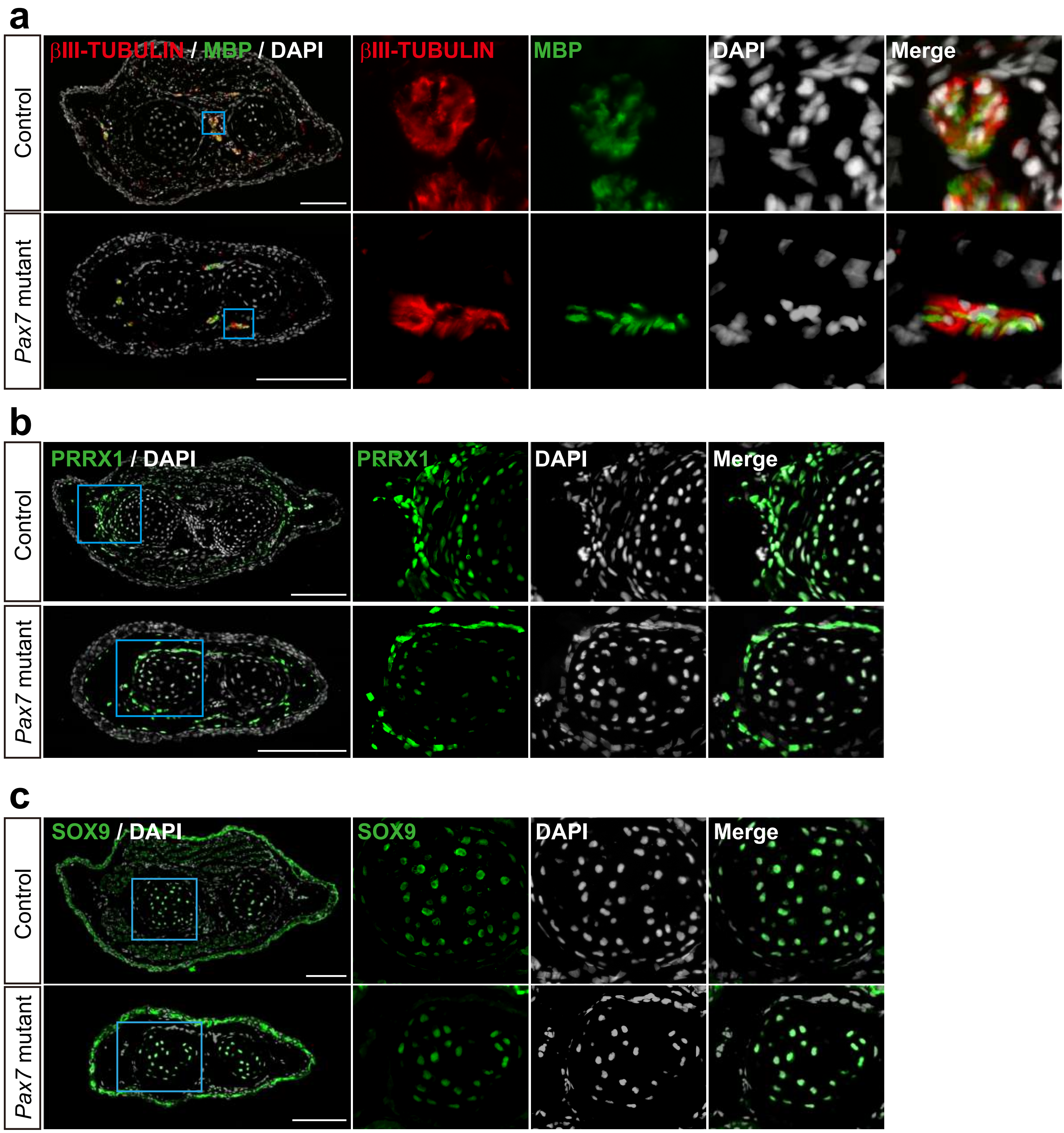
a
Control
Pax7 mutant
βIII-TUBULIN / MBP / DAPI
βIII-TUBULIN
MBP
DAPI
Merge
b
Control
Pax7 mutant
PRRX1 / DAPI
PRRX1
DAPI
Merge
c
Control
Pax7 mutant
SOX9 / DAPI
SOX9
DAPI
Merge

Fig. S2

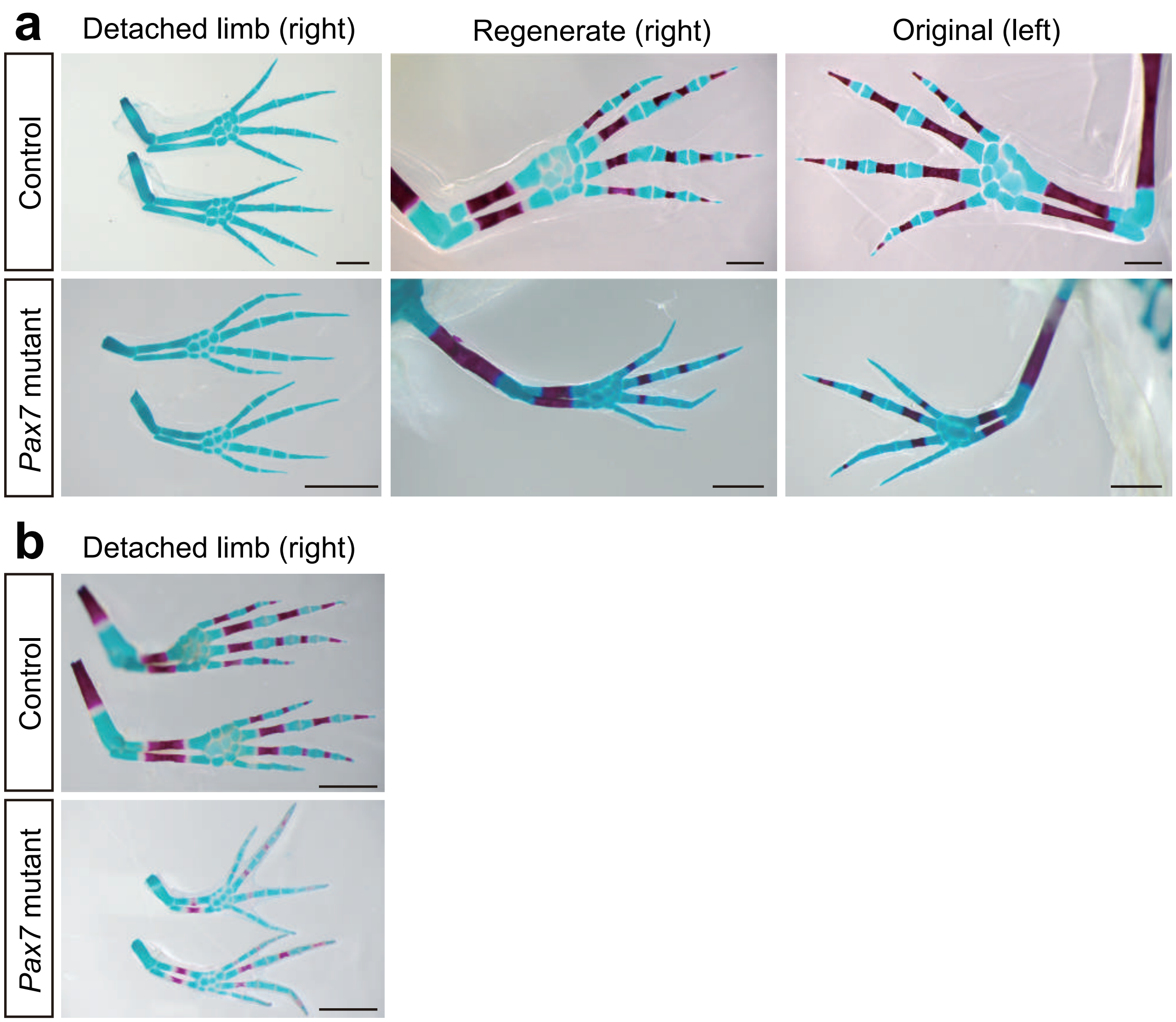

a
Detached limb (right)
Regenerate (right)
Original (left)
Control
Pax7 mutant
b
Detached limb (right)
Control
Pax7 mutant


Fig. S3

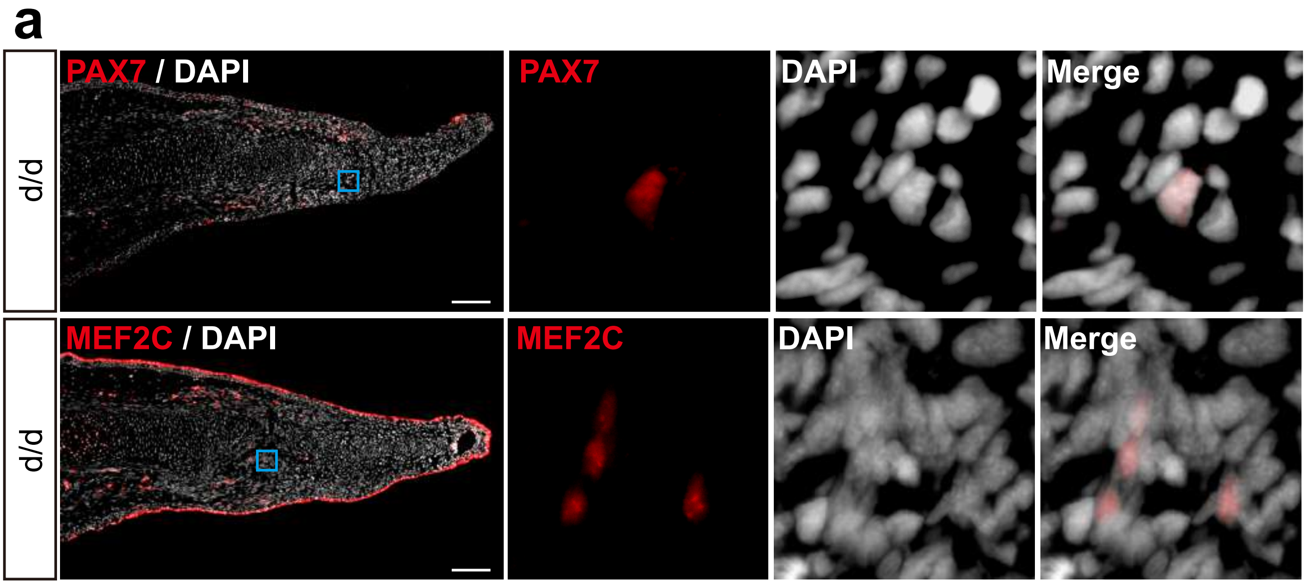


Fig. S4

### Supplementary Figure Legends

**Supplementary Fig. S1**. Lack of muscle lineage in *Pax7* mutants. (a-c) Immunofluorescence images of PAX7 (red, a), MEF2C (red, b) and MHC (red, c), and DAPI (white) in limb cross-sections, of controls and *Pax7* mutants. Boxed regions are shown at higher magnification as separated channels. Scale bars, 100 μm.

**Supplementary Fig. S2**. Examination of non-muscle lineage cell types in *Pax7* mutants. (a-c) Immunofluorescence images of βIII-TUBULIN (red), MBP (green) and DAPI (white) (a); PRRX1 (green) and DAPI (white) (b); and SOX9 (green) and DAPI (c), in limb cross-sections, of controls and *Pax7* mutants. Boxed regions are shown at higher magnification as separated channels. Scale bars, 200 μm.

**Supplementary Fig. S3**. Limb patterning in *Pax7* mutants. (a) Alcian blue and Alizarin red-stained fully regenerated (middle panels) and untouched (right panels) original forelimbs from controls (upper panels) and *Pax7* mutants (lower panels). Left panels are the stained detached right forelimbs. Amputations were carried out on 3-month-old animals, then allowed them to regenerate for 8-months for analysis. (b) The Alcian blue and Alizarin red-stained detached right forelimbs of the samples shown Fig. 1b. Scale bars, 1 mm.

**Supplementary Fig. S4**. Muscle lineage cells in 16-day regenerating blastema of d/d axolotls. (a) Immunofluorescence images of PAX7 (upper panels), MEF2C (lower panels) and DAPI (white) staining on longitudinal-sections of 16-day limb regenerates from d/d axolotls. Scale bars, 200 μm.
